# Rapid *in silico* design of antibodies targeting SARS-CoV-2 using machine learning and supercomputing

**DOI:** 10.1101/2020.04.03.024885

**Authors:** Thomas Desautels, Adam Zemla, Edmond Lau, Magdalena Franco, Daniel Faissol

**Affiliations:** Lawrence Livermore National Laboratory

## Abstract

Rapidly responding to novel pathogens, such as SARS-CoV-2, represents an extremely challenging and complex endeavor. Numerous promising therapeutic and vaccine research efforts to mitigate the catastrophic effects of COVID-19 pandemic are underway, yet an efficacious countermeasure is still not available. To support these global research efforts, we have used a novel computational pipeline combining machine learning, bioinformatics, and supercomputing to predict antibody structures capable of targeting the SARS-CoV-2 receptor binding domain (RBD). In 22 days, using just the SARS-CoV-2 sequence and previously published neutralizing antibody structures for SARS-CoV-1, we generated 20 initial antibody sequences predicted to target the SARS-CoV-2 RBD. As a first step in this process, we predicted (and publicly released) structures of the SARS-CoV-2 spike protein using homology-based structural modeling. The predicted structures proved to be accurate within the targeted RBD region when compared to experimentally derived structures published weeks later. Next we used our *in silico* design platform to iteratively propose mutations to SARS-CoV-1 neutralizing antibodies (known not to bind SARS-Cov-2) to enable and optimize binding within the RBD of SARS-CoV-2. Starting from a calculated baseline free energy of −48.1 kcal/mol (± 8.3), our 20 selected first round antibody structures are predicted to have improved interaction with the SARS-CoV-2 RBD with free energies as low as −82.0 kcal/mole. The baseline SARS-CoV-1 antibody in complex with the SARS-CoV-1 RBD has a calculated interaction energy of −52.2 kcal/mole and neutralizes the virus by preventing it from binding and entering the human ACE2 receptor. These results suggest that our predicted antibody mutants may bind the SARS-CoV-2 RBD and potentially neutralize the virus. Additionally, our selected antibody mutants score well according to multiple antibody developability metrics. These antibody designs are being expressed and experimentally tested for binding to COVID-19 viral proteins, which will provide invaluable feedback to further improve the machine learning–driven designs. This technical report is a high-level description of that effort; the Supplementary Materials includes the homology-based structural models we developed and 178,856 *in silico* free energy calculations for 89,263 mutant antibodies derived from known SARS-CoV-1 neutralizing antibodies.

## Introduction

Numerous promising therapeutic [1,2] or vaccine [3,4] research efforts are underway for COVID-19, yet a therapeutic or vaccine is not currently available. Traditional drug and biologic development are experiment-driven, typically executed via large-scale screening *in vitro* and *in vivo*. While traditional approaches have led to many effective and safe therapeutics, they require years to complete and therefore cannot alone be relied upon to support a rapid response to novel pathogens [5–7].

Data-driven (machine learning) methods are increasingly being explored for rapid therapeutic development [8,9], potentially offering the promise of a very rapid design phase. However, a purely data-driven machine learning system would be limited in rapid response scenarios due to the inevitable insufficiency of available data on novel or emerging pathogens. Theory-driven computational methods, such as molecular dynamics simulations, can provide virtually limitless amounts of data for target systems, but still require experimental validation.

Lawrence Livermore National Laboratory (LLNL) and GlaxoSmithKline (GSK) Vaccines Research have developed a combined computational-experimental platform for vaccine antigen design over the last two years. This combined platform combines experiment-driven, data-driven, and theory-driven approaches to leverage the strength of each approach, while mitigating their limitations. The initial predictions from this platform are driven by a computational component based on integrating existing experimental data, structural biology/bioinformatic modeling, and molecular dynamics simulations on high-performance computing systems. An active machine learning model aims to optimize binding behavior by iteratively proposing mutations to the amino acid sequence of an initial antigen. Proposed mutant antigens are evaluated with existing computational binding estimation tools using known or estimated antibody-antigen structures. The platform leverages a feature representation of the three-dimensional antigen-antibody interface and a Bayesian optimization algorithm to propose computational evaluation of mutants with high predicted performance and mutants that improve the machine learning model itself. The computational platform further improves its predictions by proposing designs that are evaluated with a high throughput experimental evaluation component, the results of which are incorporated into the machine learning model in a feedback loop. This combined computational-experimental antigen design platform, and results generated by it, will be described in greater detail in a future joint publication with GSK, including details of the machine learning model.

### Our approach to designing therapeutic antibodies

In response to the COVID-19 pandemic, we modified and used our computational components of the antigen design platform to propose mutations to SARS-CoV-1 neutralizing antibodies to achieve and optimize binding to the receptor binding domain (RBD) of the SARS-CoV-2 spike protein. This approach is motivated and enabled by knowledge of existing antibodies specific to the RBD of the SARS-CoV-1 spike protein that prevent SARS-CoV-1 from binding the human ACE2 receptor and entering the cell, thus neutralizing the virus [10,11]. The high similarity of SARS-CoV-1 and SARS-CoV-2, including the RBD [12], suggest that such an approach could produce efficacious therapeutic antibodies. In 22 days (1/23-2/13/2020), we demonstrated that our platform could generate antibody designs using only SARS-CoV-2 sequence information, with support from antibody/antigen structures that had previously been experimentally determined for SARS-CoV-1. We executed this entire workflow by constructing a homology-based structural model before any experimentally determined structures of the SARS-CoV-2 S-protein were available.

## Results

### Homology-based structural modeling of the SARS-CoV-2 spike protein produces accurate structural estimates when compared to experimentally derived structures

In the absence of a known SARS-CoV-2 spike protein structure, we characterized the SARS-CoV-2 surface glycoprotein sequence YP_009724390.1 [13] by constructing a homology-based structural model using the AS2TS protein modeling system [14]. The structures of the spike proteins from SARS-CoV-1 (Protein Data Bank (PDB) entries: 5×58 [15], 6nb6 [10], 2dd8 [11], and 3bgf [16]) were identified as the closest and most complete PDB structural templates to use for modeling, with resolutions 3.2, 4.2, 2.3, and 3.0 Å respectively. Of the four templates, the last three provide experimentally solved complexes with three antibodies, S230, M396, and F26G19. As our main focus in constructed homology models was to achieve the highest possible accuracy in the RBD-FAB (antigen-binding fragment) interfaces we used additional PDB templates to refine models in corresponding regions. For example, PDB template 2ghw [17] was used to improve accuracy in the modeled interface between RBD and FAB of neutralizing antibody 80R.

A CryoEM-derived structure of the SARS-CoV-2 spike protein became available on Feb. 19, 2020 and was publicly released Feb 26, 2020 [18] (https://www.rcsb.org/structure/6vsb). X-ray crystal structures (of the receptor-binding domain) became available starting March 4, 2020 (http://www.rcsb.org/structure/6VW1, to be published). Comparison with our homology models indicates that our estimated structures (completed Jan 23, 2020; publicly available beginning Feb 3, 2020) were accurate, especially at the FAB (antigen-binding fragment) antibody interfaces. **Figure 1** and **Table 1** show residue deviations for our SARS-CoV-2 homology models, SARS-CoV-2 CryoEM structures, and SARS-CoV-1 structures, using a SARS-CoV-2 X-ray structure 6w41_C as the reference. The two regions indicated by rectangular boxes (residues 382-393 and 475-484) are part of loop regions where local conformations deviate, but note that these regions are not part of the interface with the selected antibodies. **Figure 2** depicts our homology-based model superimposed on a CryoEM structure, with the M396 antibody structure provided to indicate the FAB-RBD interface region. Note that the only regions with deviations above 2.0 Å are outside the interface region.

**FIGURE 1.**
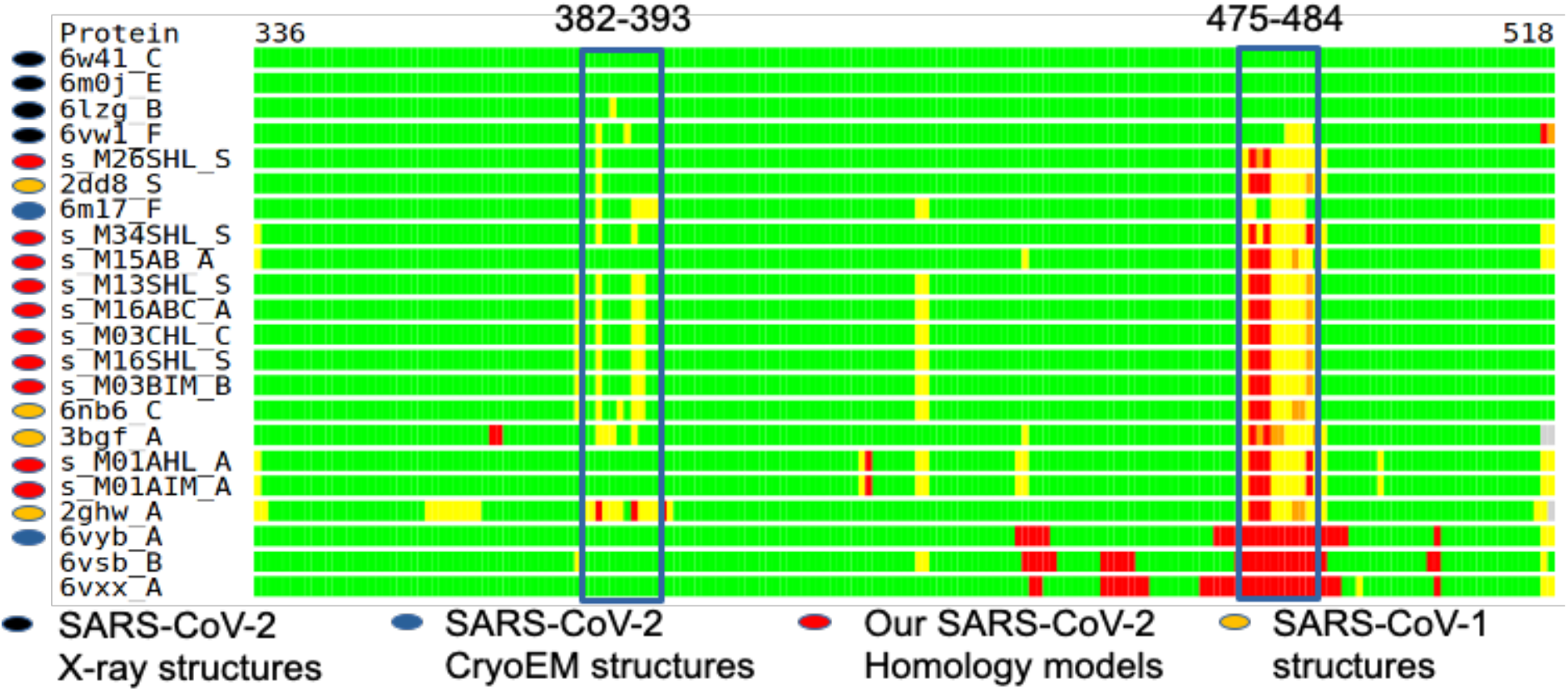
Bar representation of structural alignments with 6w41_C x-ray structure is used as reference. Four currently released X-ray structures of RBD from SARS-CoV-2 show high similarity with our SARS-C0V-2 homology models structures, SARS-CoV-2 CryoEM structures, and SARS-CoV-1 structures) in most regions except two loops (which are not part of the interface with our selected antibodies) where local conformations deviate. Green: residue deviations below 2.0 Å; yellow: below Å; orange: below 6.0 Å; red: below 8.0 Å (or not aligned/missing regions). Listed in descending order by Local-Global Alignment (LGA) [19]

**TABLE 1.**
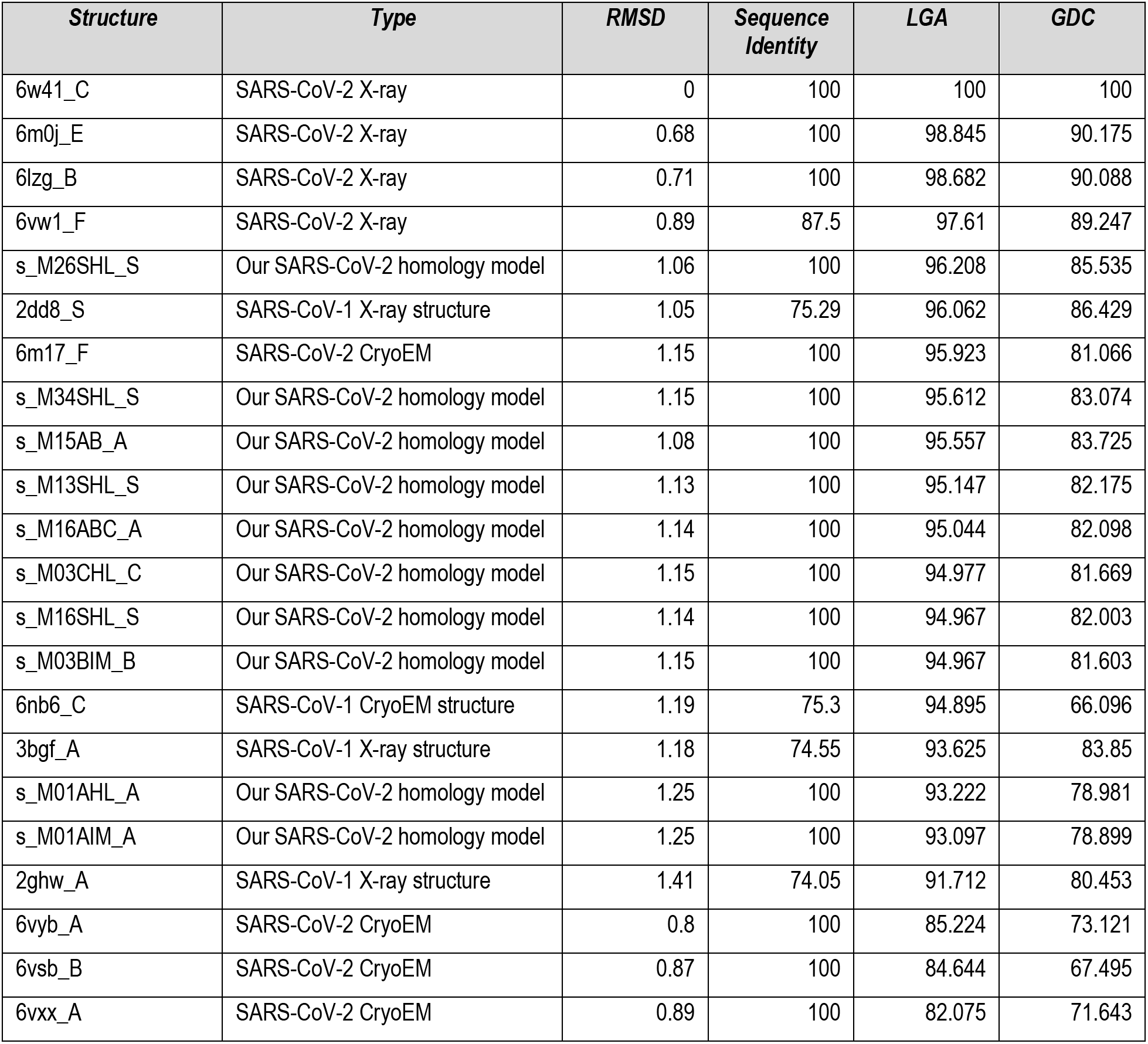
Structure similarity scores using x-ray structure 6w41 as reference. SARS-CoV-2 CryoEM structures, our SARS-CoV-2 homology models, and SARS-Cov-1 structures show similar deviations by root-mean-square deviation (RMSD), LGA [19], and global distance calculation GDC [20] from X-ray structures. Listed in descending order by LGA.

**FIGURE 2.**
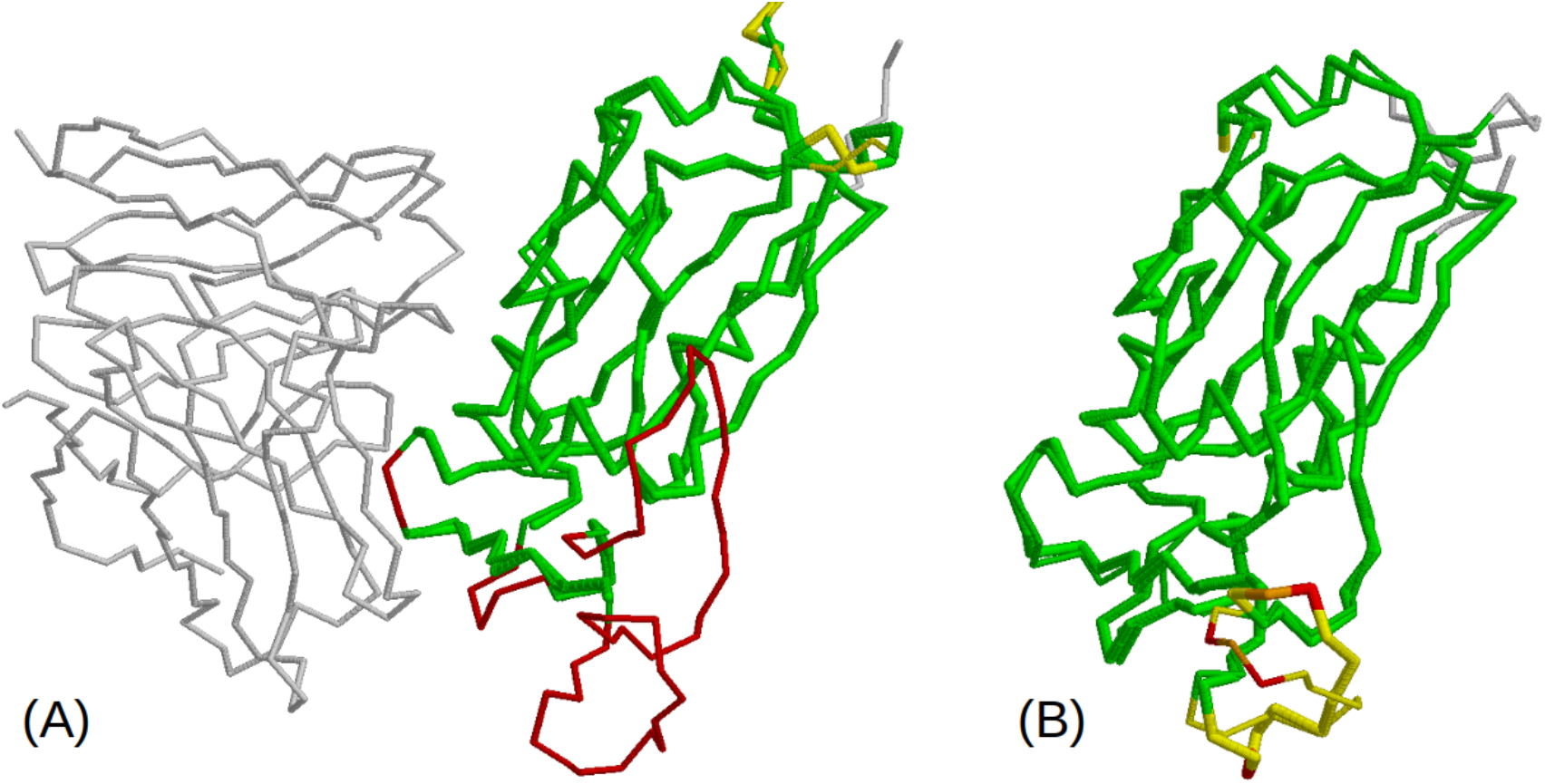
**(A)** Our homology model (thin) superimposed on CryoEM 6vsb chain A (thick). Fab (m396) structure provided to indicate FAB-RBD interface region. Regions that are missing in CryoEM structure but are present in our model are in red. **(B)** Our homology-based model superimposed with the X-ray structure 6w41_C (reference structure in Figure 1). The coloring scheme in this superposition corresponds to 5th bar in Figure 1. Regions that deviate less than 2.0 A are green. Deviations above 2.0 A are in yellow, orange or red. Note that the regions with deviations above 2.0 A are all outside the interface region

### Rapidly generating 20 initial antibody designs for experimental evaluation

Details of our computational-experimental platform will be described in greater detail in a future publication, including details on the active machine learning model, the feature space used to represent the three dimensional antigen-antibody interface, and the antibody optimization pipeline. Overall, to generate the initial 20 antibodies, we executed the following steps:

1. Obtain structures for which experimentally-determined atomic coordinates were previously deposited in the PDB for several SARS-CoV-1-neutralizing antibodies in complex with the SARS-CoV-1 S-protein; obtain sequences from SARS-CoV-2 S-protein.
2. Map public SARS-CoV-2 spike protein sequences into structures. We estimated 10 co-structures involving 3 antibodies. (Completed Jan 23, 2020; made publicly available Feb 3, 2020. We used two of these antibodies (M396 and S230) in our subsequent calculations.
3. Define a set of residues for modification in each of the starting SARS-CoV-1-neutralizing antibodies via automatic contact estimation.

- For antibody M396, up to 31 residues were allowed to simultaneously mutate, later narrowed to 21 based on intermediate results (parenthesized residues were eliminated): S31_H, (Y32_H), (T33_H), W47_H, G50_H, I51_H, (T52_H), I53_H, L54_H, I56_H, A57_H, N58_H, Y59_H, A60_H, (Q61_H), D95_H, T96_H, V97_H, (M98_H), (G99_H), G100_H, N27_L, (G29_L), S30_L, (K31_L), (W91_L), D92_L, S93_L, S94_L, D95A_L, (Y96_L). We did not consider insertions or deletions at this phase of our work.
4. Use machine learning module of computational design platform to iteratively propose mutations to the original antibody (M396) and perform free energy calculations using FoldX [21] on LLNL high-performance computing (HPC) to maximize estimated affinities to our SARS-CoV-2 spike protein homology model. By Feb 13, 2020, we evaluated 89,263 mutant antibodies selected from a design space of 10^40^ (20 amino acids^31 positions^). The primary template for this structure was PDB 2dd8.
5. Use HPC to perform additional free energy calculations on 110 selected antibody sequences by performing free energy calculations using Rosetta [22] and 102 antibodies with molecular dynamics simulations [23]; assess all sequences with the STATIUM energy prediction tool [24]. (Completed by Feb 13, 2020)
6. Assess the developability of 139 selected antibody sequences using the 5 developability metrics from the Therapeutic Antibody Profiler [25]. (Completed by Feb 13, 2020)
7. Select diverse batch of 20 M396-derived candidate antibodies for experimental based on all calculations from steps 4-6. (Completed Feb 13, 2020).

During this 22-day period, we used two high-performance computers located at LLNL to support over 200,000 CPU hours and 20,000 GPU hours, performing 178,856 in silico free energy calculations for 89,263 mutant antibodies in complex with the RBD. Given the 31 residues on M396 considered for mutation, the antibody design space has 10^40^ possibilities. Supplementary Materials include all sequences and calculations performed for M396. From these calculations, as well as from additional bioinformatic heuristics described in the Methods section, we selected 20 initial antibodies for experimental evaluation.

### Machine learning–driven computational design platform generates antibodies with improved predicted binding to SARS-CoV-2 receptor binding domain (RBD)

We used the machine learning module of our computational design platform to iteratively propose mutations to the original antibody (M396) and run FoldX [21] calculations on LLNL HPC to estimate free energies to the SARS-CoV-2 spike protein homology model. FoldX binding calculations estimate the change in free energy (ddG) of the antibody-antigen protein complex resulting from the proposed mutations (to M396). We used this calculation as part of an objective function to drive the antibody optimization via the machine learning model. We evaluated 89,263 mutants with FoldX during the course of our search for improved antibodies with estimated ddG values ranging from −10.1 to 19.2 kcal/mole. **Figure 3** plots improvements in the FoldX-estimated ddG values over the course of the machine learning-driven optimization process. These results indicate that the machine learning model was effective in searching the combinatorial space of possible antibody mutants to perform design optimization and identify increasingly improved predicted antibody designs. We then selected a subset of these antibody mutants for additional calculations using Rosetta [22], STATIUM [24], and molecular dynamics calculations [23], as well as antibody developability estimates via the Therapeutic Antibody Profiler [25] (see Methods section).

**Figure 3.**
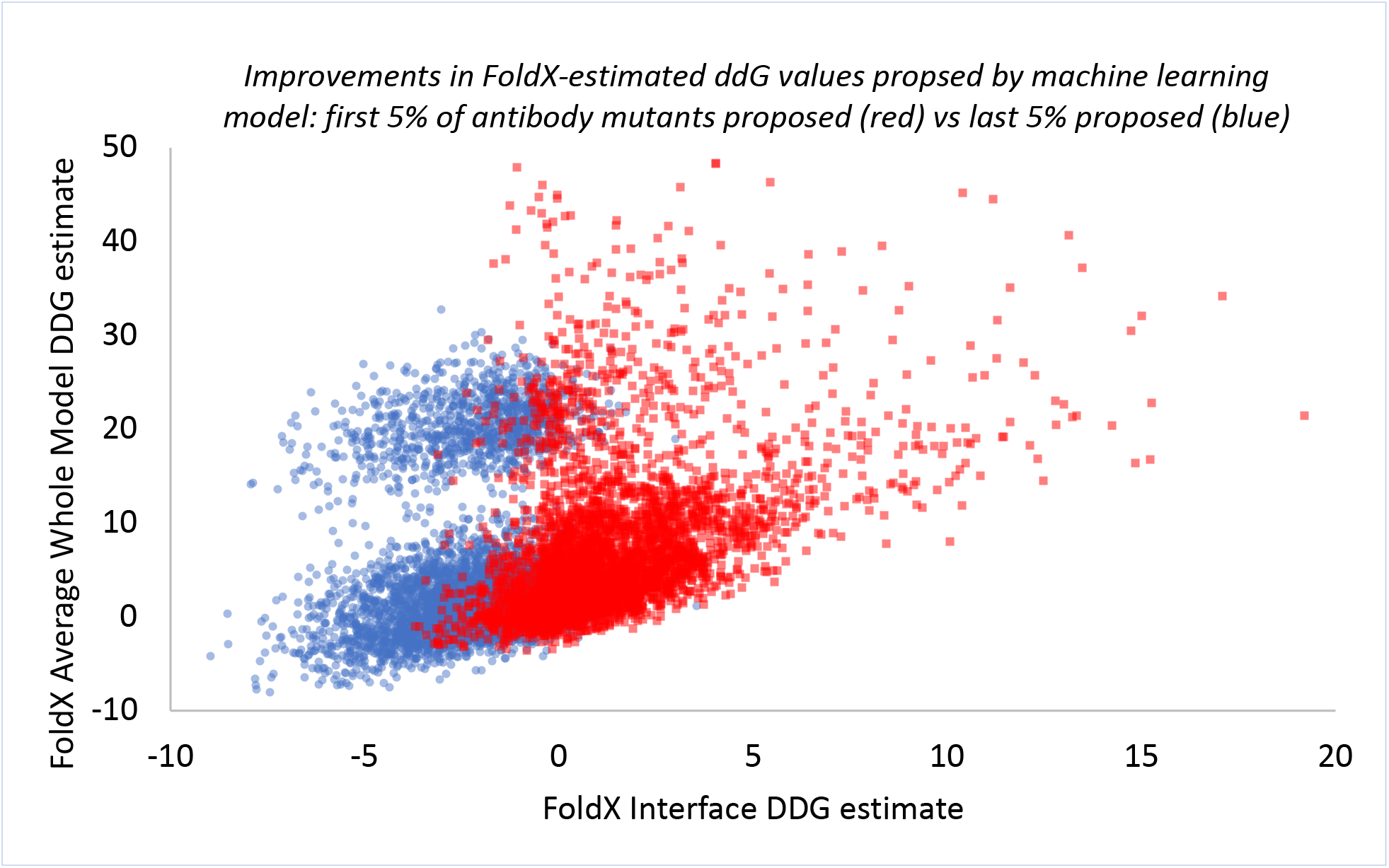
Scatter plot illustrating increasingly improved antibody mutants proposed as the machine learning-driven antibody optimization progresses. A total of 89,263 antibody mutants were proposed by the machine learning model during the course of the antibody optimization. Each antibody mutant was evaluated with FoldX with two different calculations; first to estimate energy changes for the entire complex (y-axis) and second for the RBD-FAB interface only (x-axis). The first 4,462 (5% of the total) antibody mutants proposed by the machine learning model are shown in red. The last 4,462 mutants proposed, shown in blue, resulted in a distribution of mutants with much lower energies (more favorable FAB-RDB interaction), indicating that the machine learning model was effective in searching the combinatorial space of possible antibody mutants to identify increasingly improved predicted antibody designs. Note that FoldX Interface calculations were a major driver of the objective function for optimization, and therefore show the most improvement.

While all *in silico* calculations performed were used to select our 20 first round antibodies, the molecular mechanics/generalized Born solvent accessible surface area (MM/GBSA) molecular dynamics calculations [23], described in the Methods section, are considered to be our most accurate estimate. MM/GBSA calculates antibody/antigen interaction free energies using fully solvated molecular dynamics (MD) for conformational sampling of the protein complex, but estimates free energy by a computationally less expensive implicit solvent model (GBSA). M396, our starting template for design, is known to neutralize SARS-CoV-1 by binding its RBD and preventing the virus from binding and entering the human ACE2 receptor; our MM/GBSA calculations for M396 in complex with the SARS-CoV-1 spike protein yield −52.2 kcal/mole (±7.2). M396 is known *not* to bind to the SARS-CoV-2 spike protein [18] and yields −48.1 kcal/mole (± 8.3) in complex with SARS-CoV-2 via MM/GBSA, assuming the same conformation as with SARS-CoV-1. Our 20 selected antibody designs, which are derived from M396, are predicted to have improved interaction with the RBD of the SARS-CoV-2 spike protein with MM/GBSA free energies as low as −82.0 kcal/mole, also assuming the same conformation as with SARS-CoV-1. These results suggest that these M396-derived antibody mutants potentially bind and neutralize SARS-CoV-2.

## Conclusion

Starting on Jan 23, 2020, we used two high-performance computers at LLNL over the course of 22 days to support over 200,000 CPU hours and 20,000 GPU hours, performing 178,856 *in silico* free energy calculations of candidate antibodies in complex with the SARS-CoV-2 RBD. We performed this work using just the SARS-CoV-2 sequence and previously published neutralizing antibody structures for SARS-CoV-1. Our predicted structures of SARS-CoV-2 proved to be accurate within the targeted RBD region when compared to experimentally derived structures published weeks later. With our predicted SARS-CoV-2 structures, we used our *in silico* design platform to evaluate 89,263 mutant antibodies by iteratively proposing mutations to SARS-CoV-1 neutralizing antibodies to optimize binding within the SARS-CoV-2 RBD. We selected 20 initial antibody sequences predicted to target the SARS-CoV-2 RBD. Starting from a baseline free energy of −48.1 kcal/mol (± 8.3), our 20 selected first round antibody structures are predicted to have improved interaction with the SARS-CoV-2 RBD with free energies ranging as low as −82.0 kcal/mole. The baseline SARS-CoV-1 antibody in complex with the SARS-CoV-1 RBD has a calculated free energy of −52.2 kcal/mole and neutralizes the virus by preventing it from binding and entering the human ACE2 receptor. These results suggest that our predicted antibody mutants may bind the SARS-CoV-2 RBD and therefore potentially neutralize the virus. Additionally, our antibody mutants score well according to multiple antibody developability metrics via the Therapeutic Antibody Profiler.

These 20 antibody designs are now being expressed and experimentally tested for binding to SARS-CoV-2 spike protein, which will provide invaluable feedback to further improve the machine learning–driven designs. We are also continually improving our platform and performing additional *in silico* calculations, which will be included in future updates to this document. In addition, we are currently (1) performing higher fidelity molecular dynamics calculations to increase the accuracy of predictions using [26, 27], (2) investigating binding “hotspots” via single point mutation computational analysis, and (3) exploring any potential impacts from glycosylation at the RBD site.

The Supplementary Materials to this document include (1) the 10 homology-based structural models we developed of SARS-CoV-2 in complex with SARS-CoV-1 neutralizing antibodies and (2) all antibody mutant sequences and *in silico* free energy calculations performed for 89,263 mutant antibodies derived from the M396 SARS-CoV-1 neutralizing antibody, which we used to select a first round of 20 candidate antibodies for experimental evaluation.

## Methods

### Homology-based structural modeling of the SARS-CoV-2 Spike protein structure as a starting point for antibody design

In the absence of a known SARS-CoV-2 spike protein structure, we characterized the surface glycoprotein sequence YP_009724390.1 [13] by constructing a homology-based structural model using the AS2TS system [14] and SARS-CoV-1. To assess regions of sequence-structure conservation or variability between SARS-CoV-1 and SARS-CoV-2 (sequence similarity=71%), we created a set of initial structural models using available templates. The comparative analysis between constructed models and all available templates from PDB was performed using the StralSV algorithm [28], which identifies protein fragments that exhibit structural similarities despite low primary amino acid sequence similarity and regions where some structure conformation uncertainties can be observed. We used results from these searches for modeling missing loop regions and assessment of possible structure conformation diversity in preliminary models.

In the constructed final models, the conformation of side-chain atoms was predicted using SCWRL [29] when residue-residue correspondences did not match. Residues that were identical between the template and the SARS-CoV-2 spike protein were copied from the templates onto the models. The structural and stereochemical quality of the models was checked using a contact-dot algorithm in the MolProbity software package [30], and the final constructed models were finished with relaxation using UCSF Chimera [31]. **Figures 4** **and** **5** show two of our constructed structural models of the SARS-CoV-2 spike protein in complex with a FAB (S230 and M396 in Figs. 3 and 4, respectively) of a SARS-CoV-1 neutralizing antibody. A set of all provided models was constructed to help assessment of possible different conformations in the spike-antibody complexes.

**Figure 4:**
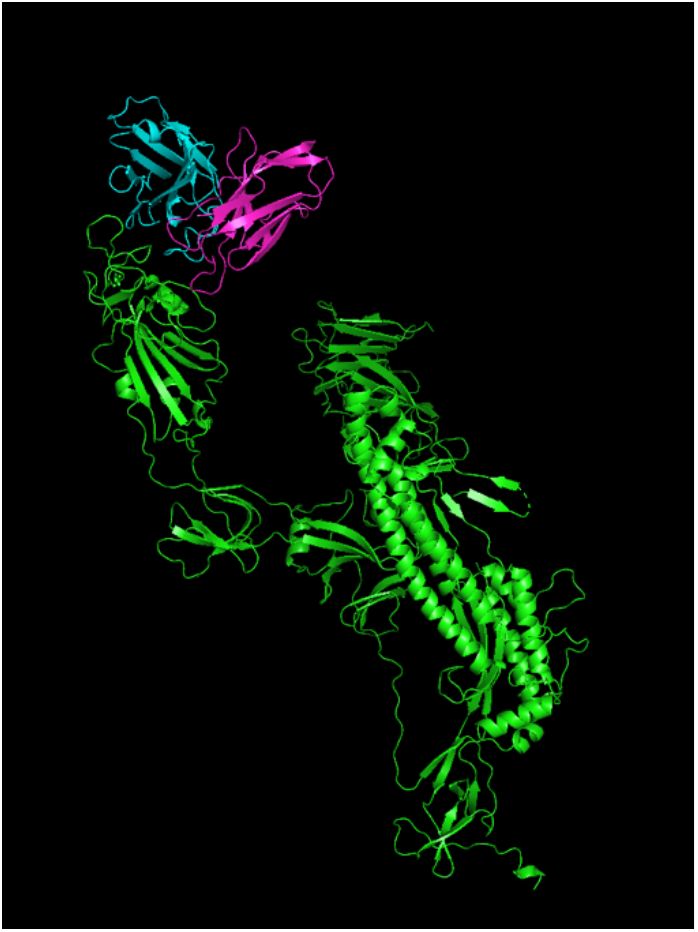
Homology-estimated structure of SARS-CoV-2 S-protein in complex with anti-SARS-CoV-1 neutralizing antibody S230 fragment: template PDB 6NB6.

**Figure 5.**
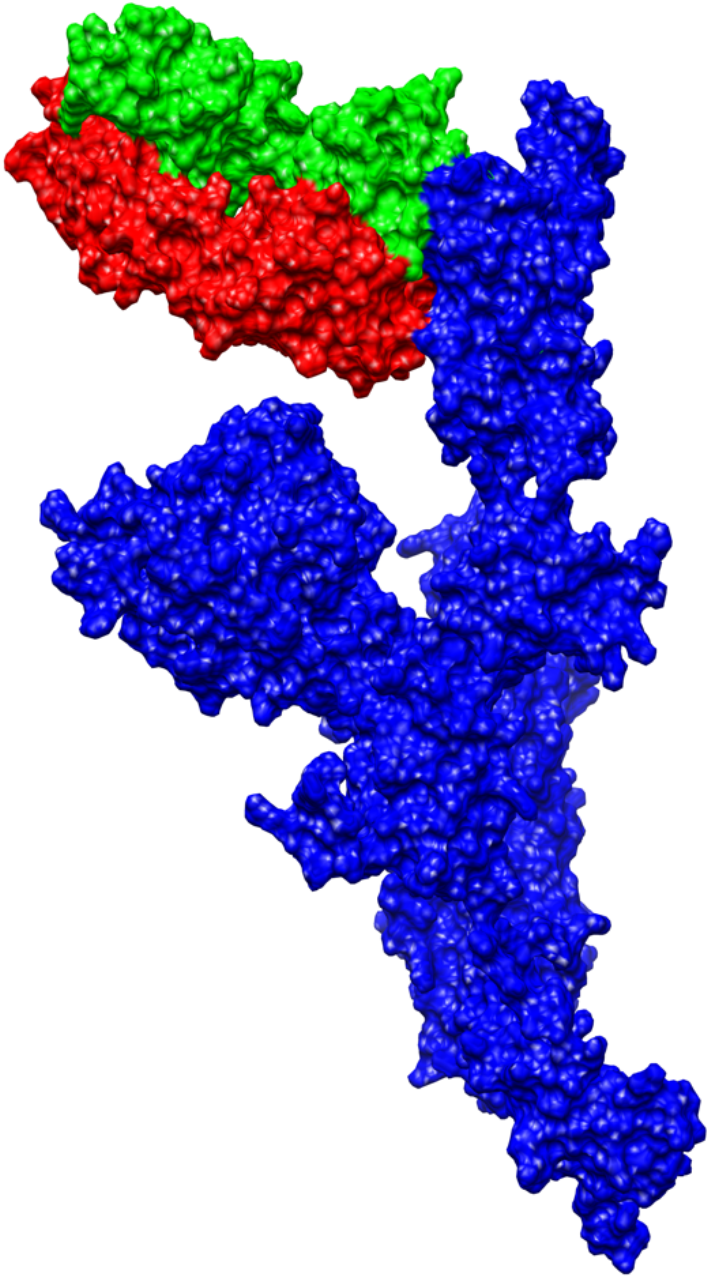
One of constructed structural models of SARS-CoV-2 Spike protein (blue) in complex with the FAB of a SARS-CoV-1 neutralizing human antibody M396 (heavy chain in green, light chain in red).

### Binding calculations with FoldX

For our ddG FoldX calculations of 89,263 mutant antibodies in complex with the RBD, we started with structure minimization using the “minimize structure” procedure available in UCSF Chimera [31]. This process was followed by up to 50 iterations of the “RepairPDB” function in FoldX. We performed ddG calculations on the defined list of mutations using the “BuildMODEL” procedure, which estimates stability (dg) defined by the free energy of a protein. We used up to 46 runs in this procedure to check if calculated rotamers of specified mutations converged to the optimal or trapped solution. A first set of our ddG estimates was calculated as a difference in free energy (dg) between the mutant and wild-type (labeled “FoldX ddG Average” in Supplementary Materials). Additional ddG estimates were calculated using the “AnalyseComplex” algorithm for which we used the wild-type and mutated models to calculate ddG based on energy changes in interface only within RBD-FAB complexes (labeled “FoldX ddG Interface” in Supplementary Materials).

### Binding calculations with Rosetta

To estimate ddG values using Rosetta, we started from structure minimization using the “relax” procedure. For each S protein-FAB complex model, we made 10 relaxed structures from which we chose the representative model with the lowest free energy. In these structures, we calculated ddG estimates as change of stability using the “ddg_monomer” algorithm with 50 iterations and settings as described in [32] (labeled “Rosetta ddG Total Energy” in Supplementary Materials). Additional ddG estimates were calculated using the “Flex_ddG” protocol [33], which evaluates energy changes by focusing on interfaces within RBD-FAB complexes only (labeled “Rosetta ddG Flex” in Supplementary Materials). In addition, Rosetta calculations (Total Energy and Flex) were performed for all possible single-point mutations for each of the 31 locations selected for mutation (19 amino acid mutation possibilities *31 locations = 589 total Rosetta single-point mutation calculations).

### Binding calculations with molecular dynamics

We used molecular mechanics/generalized Born solvent accessible surface area (MM/GBSA) calculations for antibody/antigen interaction free energies [19]. MM/GBSA uses fully solvated molecular dynamics for conformational sampling of the protein complex but estimates free energy by a computationally cheaper implicit solvent model (GBSA). All molecular dynamics simulations were performed with OpenMM [34], a toolkit for molecular simulation, using the AMBER force field [35]. Positional constraints (1 kcal/mole*Å^2^) were place on the backbone atoms (C, N, and CA) during heating of the system. The system was heated in 50 K increments for 100 ps at each temperature. Once the system was at 310 K, all constraints were removed and equilibration was performed for 9 ns. Ten individual 5 ns dynamics simulations were performed from the equilibrated system for conformational sampling. The last 1 ns of the dynamics in 50 ps increments from each of the simulations was used for the MM/GBSA calculation (200 total structures). The MM/GBSA calculations were performed using MMPBSA.py that is part of the AMBER suite of programs.

### Other decision-informing criteria

As a bioinformatic heuristic, we computed the number of mutations that are considered to be “unlikely” (labeled “number of unconventional mutations” in the dataset provided in the Supplementary Materials) for each mutant antibody considered. This heuristic describes mutations that are low probability under a mutational probability distribution derived from the Blosum62 substitution matrix [36]. This mutational probability distribution gives some substitutions higher probabilities (e.g., A->S; 0.1168) and some lower (e.g., A->W; 0.00594), where the total of the probabilities of the 19 “destination” amino acids is 1.0. If a given point mutation is given a probability of < 0.052 (i.e., less than 1/19th) the mutation is declared “unconventional” and this count is incremented by one. For example, for Alanine (A), 11 of 19 mutations are “unconventional” with this heuristic.

## Acknowledgements

The design platform leveraged in this work was supported by DARPA contract HR0011940414 and HR0011832631. We would like to thank GSK team members for their support with the design platform, including (in alphabetical order) Kathryn Arrildt, Robert van den Berg, Matthew Bottomley, Chelsy Chesterman, Jason Laliberte, Enrico Malito, Corey Mallet, Jeannette Sierra, and Newton Wahome. The COVID-19 effort described in this document was supported by Laboratory Directed Research and Development (LDRD 20-ERD-032) at Lawrence Livermore National Laboratory (LLNL). We would like to thank LLNL management, especially Shankar Sundaram for his instrumental role throughout and Tom Bates for his support with the COVID-19 effort, as well as Jason Paragas for his role in conceiving the effort.

## Supplementary Materials

Included in the Supplementary Materials are (1) the 10 homology-based structural models we developed of SARS-CoV-2 in complex with SARS-CoV-1 neutralizing antibodies and (2) all antibody mutant sequences and all *in silico* free energy calculations performed for over 89,263 mutant antibodies derived from the M396 SARS-CoV-1 neutralizing antibody used in selecting the 20 first round candidate antibodies.

This document was prepared as an account of work sponsored by an agency of the United States government. Neither the United States government nor Lawrence Livermore National Security, LLC, nor any of their employees makes any warranty, expressed or implied, or assumes any legal liability or responsibility for the accuracy, completeness, or usefulness of any information, apparatus, product, or process disclosed, or represents that its use would not infringe privately owned rights. Reference herein to any specific commercial product, process, or service by trade name, trademark, manufacturer, or otherwise does not necessarily constitute or imply its endorsement, recommendation, or favoring by the United States government or Lawrence Livermore National Security, LLC. The views and opinions of authors expressed herein do not necessarily state or reflect those of the United States government or Lawrence Livermore National Security, LLC, and shall not be used for advertising or product endorsement purposes.

