## Supplementary material for "Rapid *in silico* design of antibodies targeting SARS-CoV-2 using machine learning and supercomputing": Desautels_BioRxiv_TechReport_Coronavirus3.31.2020.pdf

---

[1] This work was performed under the auspices of the U.S. Department of Energy by Lawrence Livermore National Laboratory under contract DE-AC52-07NA27344. Lawrence Livermore National Security, LLC.

***Figure 1. Bar representation of structural alignments with 6w41_C x-ray structure is used as reference**. Four currently released X-ray structures of RBD from SARS-CoV-2 show high similarity with our SARS-C0V-2 homology models structures, SARS-CoV-2 CryoEM structures, and SARS-CoV-1 structures) in most regions except two loops (which are not part of the interface with our selected antibodies) where local conformations deviate. Green: residue deviations below 2.0 Å; yellow: below 4.0 Å; orange: below 6.0 Å; red: below 8.0 Å (or not aligned/missing regions). Listed in descending order by Local-Global Alignment (LGA) [19]*

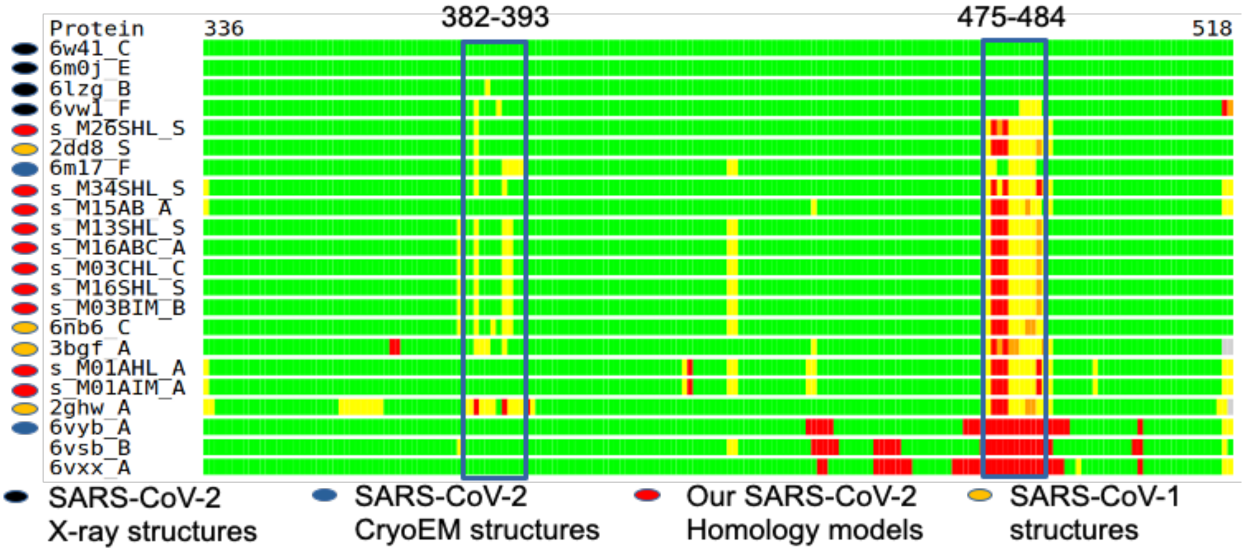


(http://www.rcsb.org/structure/6VW1 , to be published). Comparison with our homology models indicates that our estimated structures (completed Jan 23, 2020; publicly available beginning Feb 3, 2020) were accurate, especially at the FAB (antigen-binding fragment) antibody interfaces. **Figure 1** and **Table 1** show residue deviations for our SARS-CoV-2 homology models, SARS-CoV-2 CryoEM structures, and SARS-CoV-1 structures, using a SARS-CoV-2 X-ray structure 6w41_C as the reference. The two regions indicated by rectangular boxes (residues 382-393 and 475-484) are part of loop regions where local conformations deviate, but note that these regions are not part of the interface with the selected antibodies. **Figure 2** depicts our homology-based model superimposed on a CryoEM structure, with the M396 antibody structure provided to indicate the FAB-RBD interface region. Note that the only regions with deviations above 2.0 Å are outside the interface region.

***Table 1. Structure similarity scores using x-ray structure 6w41 as reference.*** *SARS-CoV-2 CryoEM structures, our SARS-CoV-2 homology models, and SARS-Cov-1 structures show similar deviations by root-mean-square deviation (RMSD), LGA [19], and global distance calculation GDC [20] from X-ray structures. Listed in descending order by LGA.*

| *Structure* | *Type* | *RMSD* | *Sequence Identity* | *LGA* | *GDC* |
|---|---|---|---|---|---|
| 6w41_C | SARS-CoV-2 X-ray | 0 | 100 | 100 | 100 |
| 6m0j_E | SARS-CoV-2 X-ray | 0.68 | 100 | 98.845 | 90.175 |
| 6lzg_B | SARS-CoV-2 X-ray | 0.71 | 100 | 98.682 | 90.088 |
| 6vw1_F | SARS-CoV-2 X-ray | 0.89 | 87.5 | 97.61 | 89.247 |
| s_M26SHL_S | Our SARS-CoV-2 homology model | 1.06 | 100 | 96.208 | 85.535 |
| 2dd8_S | SARS-CoV-1 X-ray structure | 1.05 | 75.29 | 96.062 | 86.429 |
| 6m17_F | SARS-CoV-2 CryoEM | 1.15 | 100 | 95.923 | 81.066 |
| s_M34SHL_S | Our SARS-CoV-2 homology model | 1.15 | 100 | 95.612 | 83.074 |
| s_M15AB_A | Our SARS-CoV-2 homology model | 1.08 | 100 | 95.557 | 83.725 |
| s_M13SHL_S | Our SARS-CoV-2 homology model | 1.13 | 100 | 95.147 | 82.175 |
| s_M16ABC_A | Our SARS-CoV-2 homology model | 1.14 | 100 | 95.044 | 82.098 |
| s_M03CHL_C | Our SARS-CoV-2 homology model | 1.15 | 100 | 94.977 | 81.669 |
| s_M16SHL_S | Our SARS-CoV-2 homology model | 1.14 | 100 | 94.967 | 82.003 |
| s_M03BIM_B | Our SARS-CoV-2 homology model | 1.15 | 100 | 94.967 | 81.603 |
| 6nb6_C | SARS-CoV-1 CryoEM structure | 1.19 | 75.3 | 94.895 | 66.096 |
| 3bgf_A | SARS-CoV-1 X-ray structure | 1.18 | 74.55 | 93.625 | 83.85 |
| s_M01AHL_A | Our SARS-CoV-2 homology model | 1.25 | 100 | 93.222 | 78.981 |
| s_M01AIM_A | Our SARS-CoV-2 homology model | 1.25 | 100 | 93.097 | 78.899 |
| 2ghw_A | SARS-CoV-1 X-ray structure | 1.41 | 74.05 | 91.712 | 80.453 |
| 6vyb_A | SARS-CoV-2 CryoEM | 0.8 | 100 | 85.224 | 73.121 |
| 6vsb_B | SARS-CoV-2 CryoEM | 0.87 | 100 | 84.644 | 67.495 |
| 6vxx_A | SARS-CoV-2 CryoEM | 0.89 | 100 | 82.075 | 71.643 |

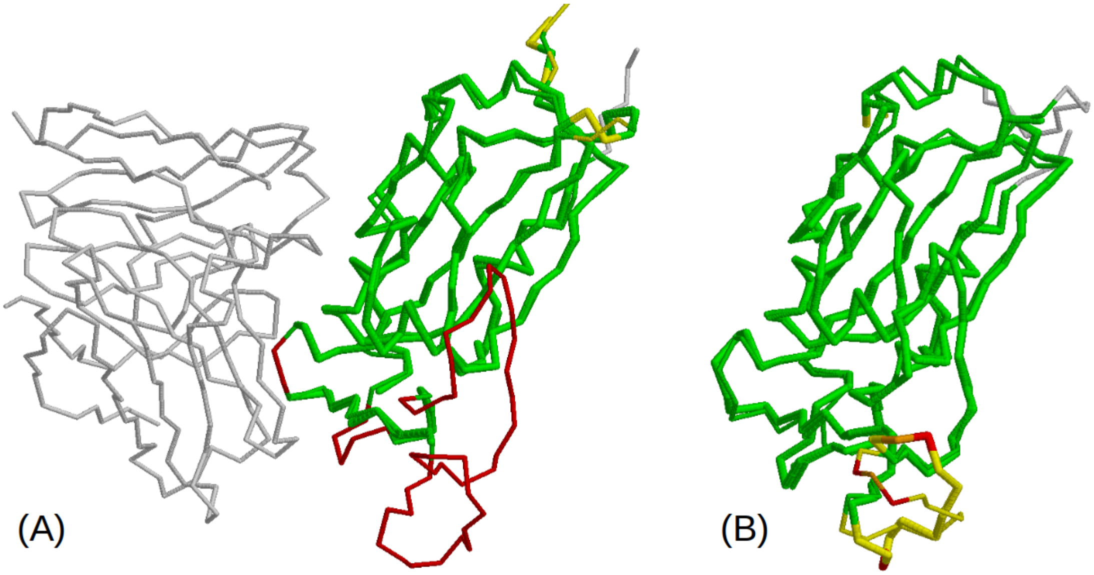


***Figure 2. (A)*** *Our homology model (thin) superimposed on CryoEM 6vsb chain A (thick). Fab (m396) structure provided to indicate FAB-RBD interface region. Regions that are missing in CryoEM structure but are present in our model are in red.* ***(B)*** *Our homology-based model superimposed with the X-ray structure 6w41_C (reference structure in Figure 1). The coloring scheme in this superposition corresponds to 5th bar in Figure 1. Regions that deviate less than 2.0 A are green. Deviations above 2.0 A are in yellow, orange or red. Note that the regions with deviations above 2.0 A are all outside the interface region*

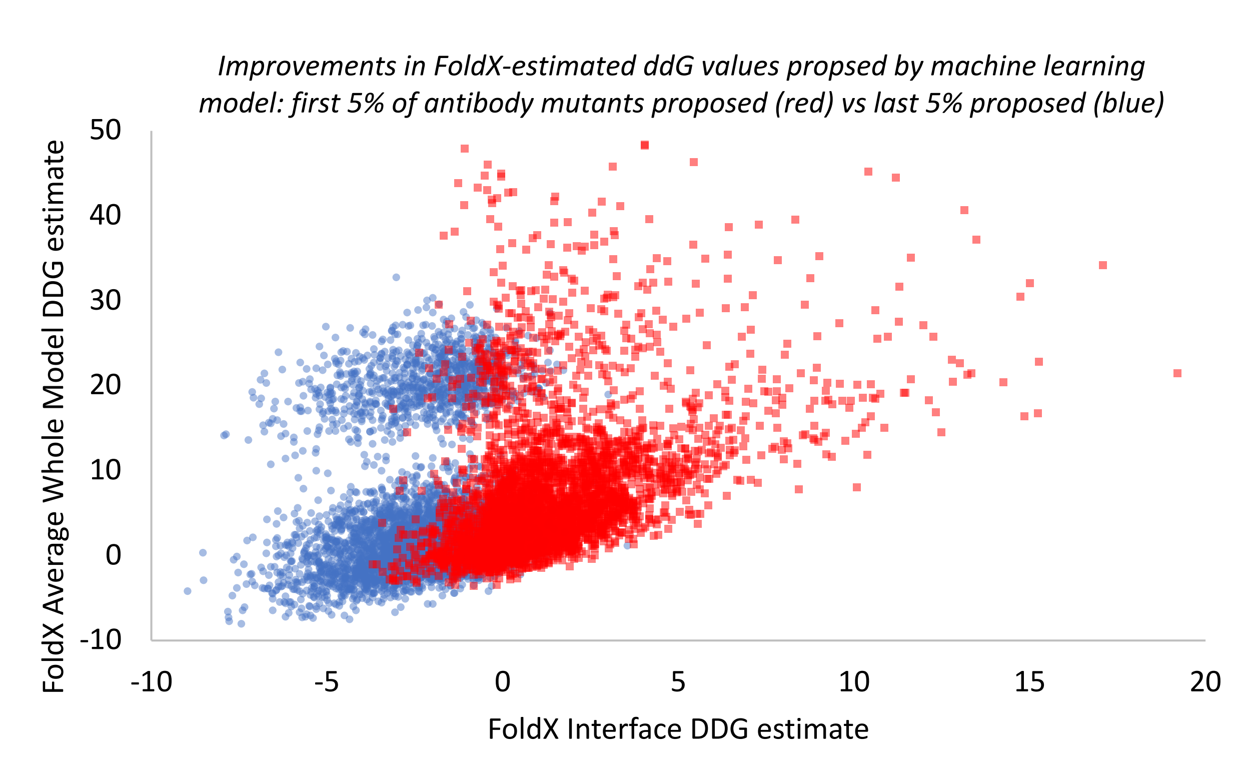


***Figure 3 Scatter plot illustrating increasingly improved antibody mutants proposed as the machine learning-driven antibody optimization progresses.*** *A total of 89,263 antibody mutants were proposed by the machine learning model during the course of the antibody optimization. Each antibody mutant was evaluated with FoldX with two different calculations; first to estimate energy changes for the entire complex (y-axis) and second for the RBD-FAB interface only (x-axis). The first 4,462 (5% of the total) antibody mutants proposed by the machine learning model are shown in red. The last 4,462 mutants proposed, shown in blue, resulted in a distribution of mutants with much lower energies (more favorable FAB-RDB interaction), indicating that the machine learning model was effective in searching the combinatorial space of possible antibody mutants to identify increasingly improved predicted antibody designs. Note that FoldX Interface calculations were a major driver of the objective function for optimization, and therefore show the most improvement.*

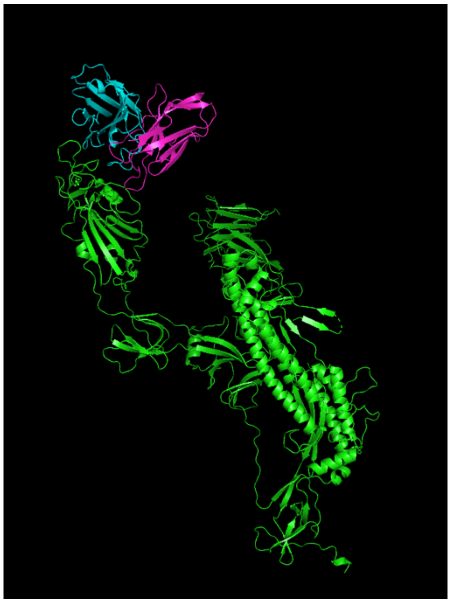

*Figure 4: Homology-estimated structure of SARS-CoV-2 S-protein in complex with anti-SARS-CoV-1 neutralizing antibody S230 fragment: template PDB 6NB6.*

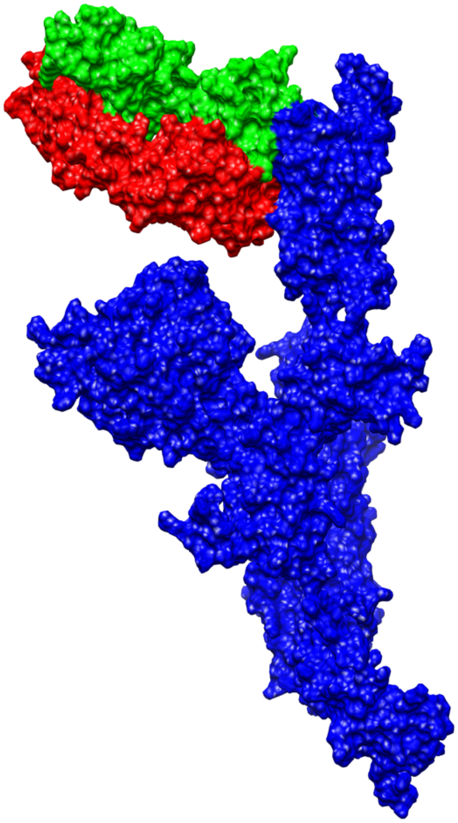

*Figure 5 One of constructed structural models of SARS-CoV-2 Spike protein (blue) in complex with the FAB of a SARS-CoV-1 neutralizing human antibody M396 (heavy chain in green, light chain in red).*
