## Supplementary material for "Rapid *in silico* design of antibodies targeting SARS-CoV-2 using machine learning and supercomputing": README.docx

In response to COVID-19, we used our in-silico design platform to propose mutations to SARS-CoV-1 neutralizing antibodies to achieve and optimize binding to the receptor binding domain of SARS-CoV-2 spike protein. The data contained here is a standard .csv file. Each row represents a unique mutant antibody proposed by our machine learning model, derived from a publicly available starting template antibody M396. The data for each mutant antibody (i.e., each row) are along the columns and are the result of several software-based calculations as described in the main text and below. Fields (column headers), in the order they appear in the file, are:

- Antibody_ID
  - *unique antibody identifier*

- Complex

- *an identifier for the protein structure used for software calculations also included in Supplementary Materials*
- Structure_MD5_Hash
  - *MD5 hash protein structure used for software calculations*
- Antibody
  - *identifier for the starting template antibody from which the mutant antibody was derived*
- Mutation
  - *the mutations performed to produce the mutant antibody from the starting template antibody*
  - *see conversion between our internal sequence ordering and original ordering)*
- Antibody_Sequence
  - *sequence of the mutant antibody for which calculations were performed*
- FoldX_Average_Whole_Model_DDG
  - *Antibody-antigen free energy estimates using FoldX [1] software for the mutant antibody for the whole Antibody-antigen complex*
- FoldX_Average_Interface_Only_DDG
  - *Antibody-antigen free energy estimates using FoldX [1] software for the mutant antibody for the interface of the Antibody-antigen complex*
- Rosetta_Flex_DDG
  - *Antibody-antigen bindings estimates using Rosetta [2] software for the mutant antibody using the "Flex_ddG" protocol*
- Rosetta_Total_Energy_DDG
  - *Antibody-antigen bindings estimates using Rosetta [2] software for the mutant antibody using the “ddg_monomer" algorithm*
- MMGBSA
  - *Antibody-antigen bindings estimates from LLNL molecular dynamics calculations (molecular mechanics/generalized Born solvent accessible surface area) for the mutant antibody*
- Statium
  - *Antibody-antigen predictions using Statium [3] software for the mutant antibody*
- Total_CDR_Length
  - *Total CDR Length from Therapeutic Antibody Profiler [4]*
- CDR_Vicinity_PSH_Score
  - *CDR Vicinity PSH Score (Kyte & Doolittle) from Therapeutic Antibody Profiler [4]*
- CDR_Vicinity_PPC_Score
  - *CDR Vicinity PPC Score from Therapeutic Antibody Profiler [4]*
- CDR_Vicinity_PNC_Score
  - *CDR Vicinity PNC Score from Therapeutic Antibody Profiler [4]*
- SFvCSP_Score
  - *SFvCSP Score from Therapeutic Antibody Profiler [4]*
- number_of_unconventional_mutations
  - *A bioinformatic heuristic representing how evolutionarily unlikely the mutations in the mutant antibody are. Derived from BLOSUM62 [5]*
- Sum_of_Rosetta_Flex_single_point_mutations
  - *Sum of Rosetta Flex calculations for each single point mutations present in mutant*
- Sum_of_Rosetta_Total_Energy_single_point_mutations
  - *Sum of Rosetta Total Energy calculations for each single point mutations present in mutant*
- Rosetta_Flex_calculations_single_point_mutations
  - *Rosetta Flex calculations of single point mutations for each mutation present in the mutant antibody*
  - *ordering corresponds to ordering in field labeled “Residue_locations_allowed_to_mutate_LLNL”*
  - *Note not all locations are mutated for every mutant.*
- Rosetta_Total_Energy_calculations_single_point_mutations
  - *Rosetta Total Energy calculations of single point mutations for each mutation present in the mutant antibody.*
  - *ordering corresponds to ordering in field labeled “Residue_locations_allowed_to_mutate_LLNL”*
  - *Note not all locations are mutated for every mutant.*
- Residue_locations_allowed_to_mutate_LLNL
  - *Residue locations allowed to mutate using our internal LLNL numbering of sequences. Note not all locations are mutated for every mutant*
- Residue_locations_allowed_to_mutate_original
  - *Residue locations allowed to mutate using original numbering of sequences. Note not all locations are mutated for every mutant.*

Note that the residue numbering in our structures differs from the numbering in the original M396 structure. The conversion is as follows:

| Our numbering | Original M396 numbering |
| --- | --- |
| S31 | S31_H |
| Y32 | Y32_H |
| T33 | T33_H |
| W47 | W47_H |
| G50 | G50_H |
| I51 | I51_H |
| T52 | T52_H |
| I54 | I53_H |
| L55 | L54_H |
| I57 | I56_H |
| A58 | A57_H |
| N59 | N58_H |
| Y60 | Y59_H |
| A61 | A60_H |
| Q62 | Q61_H |
| D99 | D95_H |
| T100 | T96_H |
| V101 | V97_H |
| M102 | M98_H |
| G103 | G99_H |
| G104 | G100_H |
| N271 | N27_L |
| G273 | G29_L |
| S274 | S30_L |
| K275 | K31_L |
| W335 | W91_L |
| D336 | D92_L |
| S337 | S93_L |
| S338 | S94_L |
| D340 | D95A_L |
| Y341 | Y96_L |

[1] Schymkowitz J, et al. The FoldX web server: an online force field. Nucleic Acids Res, 2005 Jul 1, Volume 33, Issue Web Server issue, p.W382-8

[2] Leaver-Fay A, et al. ROSETTA3: an object-oriented software suite for the simulation and design of macromolecules. Methods in Enzymology, 2011; Vol. 487, p 545-574.

[3] DeBartolo J, et al. Genome-wide prediction and validation of peptides that bind human prosurvival Bcl-2 proteins. PLoS computational biology. 2014 Jun 26;10(6):e1003693.

[4] Raybould, M. I., et al. Five computational developability guidelines for therapeutic antibody profiling. Proceedings of the National Academy of Sciences, 116(10), 4025-4030 (2019).

[5] Henikoff, S., & Henikoff, J. G. (1992). Amino acid substitution matrices from protein blocks. Proceedings of the National Academy of Sciences, 89(22), 10915-10919.
